# The effects of brief heat during early booting on reproductive, developmental and physiological performance in common wheat (*Triticum aestivum* L.)

**DOI:** 10.1101/2022.02.20.481180

**Authors:** Jiemeng Xu, Claudia Lowe, Sergio G. Hernandez-Leon, Susanne Dreisigacker, Matthew P. Reynolds, Elisa M. Valenzuela-Soto, Matthew J. Paul, Sigrid Heuer

## Abstract

Rising temperatures due to climate change threaten agricultural crop productivity. As a cool-season crop wheat is heat sensitive, but often exposed to high temperatures during cultivation. In the current study, a bread wheat panel of spring wheat genotypes, including putatively heat-tolerant Australian and CIMMYT genotypes, was exposed to a 5-day mild (34°C/28°C, day/night) or extreme (37°C/27°C) heat stress during the sensitive pollen developmental stage. Worsening effects on anther morphology were observed as heat stress increased from mild to extreme. Even under mild heat a significant decrease in pollen viability and grain number per spike from primary spike was observed compared with the control (21°C/15°C), with Sunstar and two CIMMYT breeding lines performing well. A heat-specific positive correlation between the two traits indicates the important role of pollen fertility for grain setting. Interestingly, both mild and extreme heat induced development of new tillers after the heat stress, providing an alternative sink for accumulated photosynthates and significantly contributing to the final yield. Measurements of flag leaf maximum potential quantum efficiency of Photosystem II (Fv/Fm) showed an initial inhibition after the heat treatment, followed by a full recovery within a few days. Despite this, model fitting using chlorophyll SPAD measurements showed an earlier onset or faster senescence rate under heat stress. The data presented here provide interesting entry points for further research into pollen fertility, tillering dynamics and leaf senescence under heat. The identified tolerant wheat genotypes can be used to dissect the underlying mechanisms and breed climate-resilient wheat.

## INTRODUCTION

Wheat is one of the most important crops for human consumption, grown on 220 million hectares with a total production of 760 million tons in 2020 (FAOSTAT). In climate change scenarios, wheat plants are prone to be exposed to warmer and more variable temperatures (Trnka et al. 2014). Beyond a physiological threshold, high temperatures cause stress and impair plant growth and development. Both historical data and future predictions have revealed the negative effects of heat on wheat productivity at the global and regional scale (Liu et al. 2016; Zampieri et al. 2017; Pequeno et al. 2021). Therefore, it is crucial to identify and breed heat-adapted varieties to sustain wheat production and ensure food security.

In nature, the adverse effects of heat stress on plants can be variable depending on the intensity, duration and developmental stage (Yeh et al. 2012). Most heat-related studies in wheat have been field based and used late sowing to expose plants to high temperatures during the flowering and grain filling stages, however, short episodes of heat during earlier reproductive stages can also cause significant damage (Zampieri et al. 2017). Indeed, anther and pollen development are considered to be the stages most vulnerable to heat stress (Zinn et al. 2010; Rieu et al. 2017). Stage-specific treatments have found that wheat is particularly sensitive to heat around eight days before anthesis, which coincides with the early meiosis to tetrad stage of pollen development (Saini and Aspinall 1982; Prasad and Djanaguiraman 2014). Because pollen development occurs during the booting stage, while spikes are still inside the developing pseudostem in wheat, the length of the auricles between the flag leaf and the penultimate leaf (referred as auricle interval length, AIL) has been used as a proxy for pollen development. AIL of between 3-6 cm has been associated with this sensitive stage (Erena et al. 2021; Bokshi et al. 2021). Brief heat exposure during this sensitive period resulted in abnormal meiosis behavior (Omidi et al. 2014; Draeger and Moore 2017) and significant reduction in pollen fertility (Prasad and Djanaguiraman 2014; Begcy et al. 2018; Browne et al. 2021). Because booting usually occurs during the cooler part of the cropping season, and because of the difficulties in applying precise stage-specific heat stress, few studies have examined the natural variation in pollen viability under heat stress and its association with yield (Bheemanahalli et al. 2019; Bokshi et al. 2021). However, considering a warmer and increasingly erratic climate, this area warrants further investigation.

Spike number is one of the main components determining wheat yield; it is very variable and responsive to environmental factors (Slafer et al. 2014). Interestingly, contrasting responses of spike number under heat stress have been reported. When exposed to continuous high temperatures during the terminal flowering and grain filling stages, spike formation and tillering were always reduced (Cai et al. 2016; Sharma et al. 2016; Dwivedi et al. 2017; Kumar et al. 2021). In contrast, after a short episode of heat stress during earlier developmental stages, spike numbers increased (Bányai et al. 2014; Hütsch et al. 2019; Chavan et al. 2019). Enhanced spike formation after early heat stress is surprising, but the underlying tillering dynamics and impact on final yield have not been dissected. In addition to the effects on pollen fertility and spike number, heat-induced yield loss has also been ascribed to accelerated leaf senescence, shortening the duration for grain filling (Cossani and Reynolds 2012; Shirdelmoghanloo et al. 2016; Pinto et al. 2016; Sade et al. 2018; Bergkamp et al. 2018). As indicators of senescence, chlorophyll SPAD (chlorophyll content index) (Richardson et al. 2002) and Fv/Fm (the maximum potential quantum yield of photosystem II) (Murchie and Lawson 2013) have been widely used to evaluate this trait. Under terminal heat, SPAD and Fv/Fm were often reduced in leaf tissue during senescence and closely related to yield contributing traits, such as thousand grain weight (Talukder et al. 2014; Hassan et al. 2018; Mirosavljević et al. 2021; Touzy et al. 2022). Studies for the genetic analysis of these leaf senescence related traits are also available (Azam et al. 2015; Bhusal et al. 2018; Touzy et al. 2022). Nevertheless, time course measurements of SPAD and Fv/Fm, which enable model fitting and senescence parameter prediction, have rarely been captured in wheat under heat stress (Pinto et al. 2016; Šebela et al. 2020; Touzy et al. 2022), especially after brief heat during the early reproductive stage. In the present study, a wheat heat panel, including putatively heat-tolerant Australian and CIMMYT-nominated spring wheat genotypes, was exposed to a 5-day heat stress during early pollen developmental stage and analyzed for effects on (i) pollen viability and seed set; (ii) spike formation and underlying tillering dynamics; (iii) leaf senescence measured with SPAD and Fv/Fm; and (iv) relationships among the reproductive, developmental, physiological and yield-related traits.

## MATERIALS AND METHODS

### Plant materials

In this study, three greenhouse experiments (named Exp1, Exp2 and Exp3) were conducted. For Exp1 and Exp2, the same set of wheat 14 genotypes were used, and 22 lines (7 overlapping with Exp1 and Exp2) were grown in Exp3 (See Table S1 for genotype details). These lines are putatively heat-tolerant elite spring varieties (Sunstar, Sokoll and Waagaan), parental lines and their pre-breeding materials (Cossani and Reynolds 2015; Erena 2018), as well as two lines from the United Kingdom included as controls.

### Plant cultivation and heat stress treatment

The plants for all the experiments were grown in the controlled environment and glasshouse facilities at Rothamsted Research, Harpenden, UK (51.8094° N, 0.3561° W). All experiments followed a split randomized complete block design (split RCBD) with 4 blocks/biological replicates. Different genotypes were assigned randomly to the whole plot within each block, and temperature treatments (control/CT and heat/HT) were then assigned to the subplots within each whole plot. Two seeds were sown in separate pots filled with Rothamsted Standard compost (75% medium grade peat; 12% screened sterilized loam; 3% medium grade vermiculite; 10% 5mm screened lime free grit) and fertilized with Osmocote Exact 3-4 month at the rate of 3.5kg m^-3^. One week after sowing, seedlings were thinned down to 1 per pot and grown under natural light glasshouse conditions with 16-h light period supplemented with artificial light (230W LED, Kroptek Ltd., London, UK) if natural light intensity fell below 175 µmol m^−2^ s^−1^. The temperature in the glasshouse was set at 21°C/15°C (day/night, actual value: 21.5 ± 0.4/16.3 ± 0.5°C for Exp1; 20.6 ± 0.8/15.4 ± 0.7°C for Exp2; 21.7 ± 0.4/15.4 ± 0.3°C for Exp3) and the relative humidity (RH) was around 60/75% (day/night) for Exp1, 57/69% for Exp2 and 42/49% for Exp3 (Fig S1 a, b, c). At the booting stage, plants where the primary tiller reached the targeted auricle interval length (AIL, 6 cm for Exp1, 2-3 cm for Exp2 and Exp3, see actual value of AIL in Fig S2) were sequentially moved into Fitotron® Modular Plant Growth Chambers (HGC1514, Weiss Technik UK Ltd., Loughborough, UK) for HT treatment (Exp1: 36.97 ± 0.03/26.95 ± 0.17°C, Exp2: 37.02 ± 0.01/27.00 ± 0.01°C, Exp3: 34.00 ± 0.02/28.00 ± 0.01°C). The light period was 16 hours and the intensity was maintained around 600 µmol m^−2^ s^−1^ at plant level. The RH was maintained between 70-75% (Fig S1 d, e, f). In Exp1, plants for CT treatment were kept in the glasshouse, while in Exp2 and Exp3, similar growth chambers as heat stress were used for CT plants (Exp2: 21.01 ± 0.01/15.01 ± 0.01°C, Exp3: 21.01 ± 0.02/15.01 ± 0.01°C; light period and intensity and RH was the same as heat treatment (Fig S1 e, f). Plants for both CT and HT treatments were kept in the corresponding conditions for 5 days and then moved back to the glasshouse until final harvest.

### Morphological, phenological and chlorophyll physiological measurements

On the day before (day 0) and after (day 6) HT treatment, the primary tiller of each plant was tagged and measured for AIL. Plant height was also recorded at these two time points in Exp2 and Exp3 (Fig S1 g). Chlorophyll SPAD and Fv/Fm (maximum potential quantum efficiency of Photosystem II) were measured at the same time as AIL and weekly thereafter (Fig S1 g). The SPAD measurement was performed with a MC-100 Chlorophyll Concentration Meter (Apogee Instruments, Inc., Logan, UT, USA). Fv/Fm was measured with a Pocket PEA (Hansatech Instruments Ltd., Norfolk, UK) after 15-20 min dark adaptation. For each plant, the mean SPAD value of measurements at the tip, middle and bottom of flag leaf was obtained and one measurement of Fv/Fm was made in the middle of flag leaf. After the HT treatment, the heading date of each plant was recorded to calculate days to heading in Exp2 and Exp3. Physiological maturity of the spike on the tagged tiller was recorded as days to maturation.

### Measurement with tagged tillers/spikes for pollen fertility and grain number per spike

During anthesis in Exp2 and Exp3, the 4^th^ or 5^th^ spikelet (counted from the bottom) was sampled from the tagged tiller. One anther from the bottom two florets was photographed for a representative image and measured for anther length. In Exp3, the remaining five anthers from the bottom two florets were pooled together for pollen viability analysis using staining with Lugol’s solution. Fully stained pollen was scored as viable, whereas partially stained or aberrant shape pollen were scored as non-viable. At maturity, the number of filled grains of the tagged spike was counted and recorded as grain number per spike, and spike length and spikelet number were measured.

### Measurement of tillering dynamic change

In Exp1, development of extra young spikes after heat stress was observed (Fig 2 a). In Exp2 and Exp3, tiller number was therefore continuously counted for CT and HT treated plants on the day (day 0) before and after (day 6) the 5-day HT treatment, and on weekly intervals thereafter until maximum tillering (Fig S1 g).

### Yield related measurements at maturation

At maturity, the spikes per plant were distinguished into “old” spikes (labelled just before starting the HT treatment in Exp2 and Exp3) and “new” spikes (Fig 2 a), harvested separately, then oven dried at 40°C for seven days prior to mechanical threshing and cleaning. The weight, number, length and width were then determined for grain samples from old and new spikes separately with a scale and a MARViN digital seed analyser (MARViTECH GmbH., Wittenburg, Germany). Grain yield per plant was calculated as the sum of grains from old and new spikes. The aboveground biomass for each plant was determined as the dry weight of all straw materials dried in an oven at 80°C for forty-eight hours.

### Statistical analysis

The data from the time course SPAD measurements were fitted using a generalized additive model (GAM) for each of the three experiments to estimate maximum SPAD (SPADmax), senescence onset (SenOnset) and senescence rate (SenRate) (graphic illustration in Fig S3). SPAD was predicted by a smooth function of time (days counted from stress initiation), with a separate smooth function fitted for each combination of genotype and treatment. The Exp1 model used eight basis functions, whereas Exp2 and Exp3 used seven. SPADmax was estimated from the fitted predicted model. SenOnset was calculated as the day that SPAD fell to 95% of the maximum SPAD. Senescence period was defined over fourteen days from the onset or until the end of the measurement, whichever was shorter. SenRate was then calculated as the daily reduction of SPAD over the senescence period. GAMs were fitted in R (version 3.6.1) using the “mgcv” package (version 1.8-35) (Wood, 2011).

All trait measurements and calculated parameters (see Table S2 for details) were used for statistical analysis in R 4.0.3 (https://www.R-project.org/). Firstly, descriptive statistics were summarized with the “describeBy” from the “psych” package. The effects of genotype treatment and the interaction were obtained from ANOVA analysis with the model fitted with “lmer” from the R package “lmerTest”. In the model, genotype, treatment and the interaction were treated as fixed factors, while block and genotype nested in block as random effects. Afterwards, Tukey’s post-hoc test was carried out for multiple test comparison to identify genotypic variation. Estimated marginal means were calculated for each combination of genotype and treatment. Subsequently, for either CT or HT treatment, Pearson correlation coefficient table was calculated by using “tab_corr” from “sjPlot” package among measurements and pairwise-deletion method was used to account for missing data. For each experiment and temperature treatment, correlations among different traits were visualized as networks with the “qgraph” package.

## RESULTS

### Heat impaired pollen fertility and grain number per spike

To understand the effects of heat on pollen development and grain setting, the primary tiller of each plant was tagged and measured. In Exp1 and Exp2, the imposed severe heat treatment of 37°C/27°C caused nearly complete loss of grain setting for all genotypes, except for Paragon and Cadenza (Fig S4). The anther morphology was also severely changed by the HT treatments indicating complete absence of viable pollen (Fig 1 a, b). In Exp3, relatively mild heat stress (34°C/28°C) also significantly reduced anther length, however, this was less severe compared to Exp2 (Fig 1 a, b) and pollen viability was therefore analyzed with Lugol’s solution staining. The results showed considerable variation among genotypes, ranging from 0% to 60%. One line (SWBL1.1, a progeny between the cross of Sokoll and Weebill1) had the highest pollen viability (relative to control value), followed by SWES, SUN (Sunstar) and WBL1.2 (Weebill1) (Fig 1 c). Grain number of the tagged primary spike was also variable among the genotypes with SUN showing the highest relative to control value (Fig 1 d). Further analysis found a positive correlation between pollen viability and grain number per spike under HT (Fig 1 f), but not under CT treatment (Fig 1 e).

**Fig 1.**
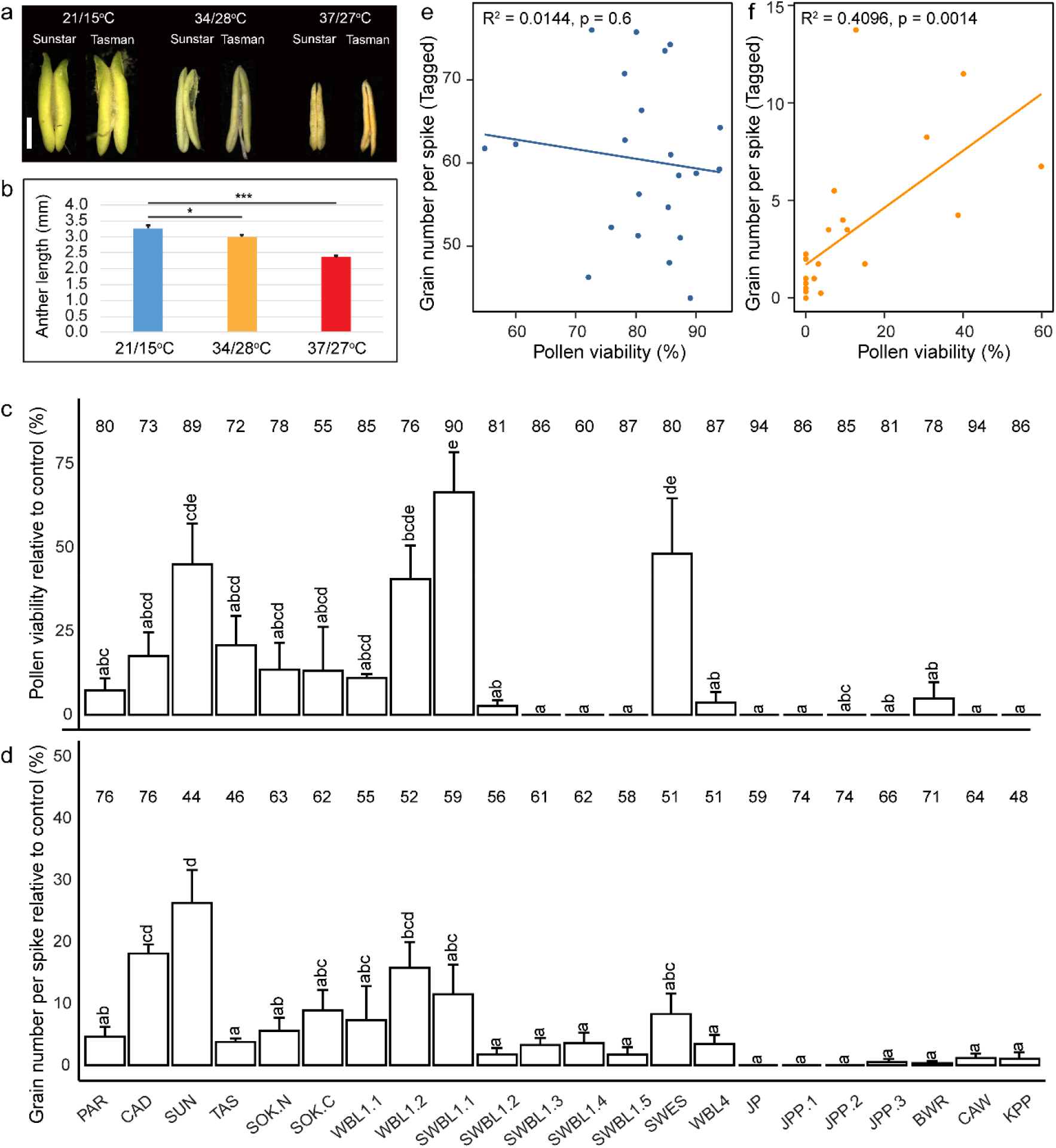
The heat effects on wheat reproductive traits. Anther morphology after heat treatments in two representative genotypes is shown in (a). Average anther length across all analyzed genotypes in Exp2 (37°C/27°C) and Exp3 (34°C/28°C) is shown in (b). Genotypic variation of pollen viability (c) and grain number per spike (d) under heat in Exp3. The data were obtained from the primary tiller and calculated as relative to control values, which are shown on top of each bar. Pearson correlation between pollen viability and grain number per spike under control (e) and heat (f).

**Fig 2.**
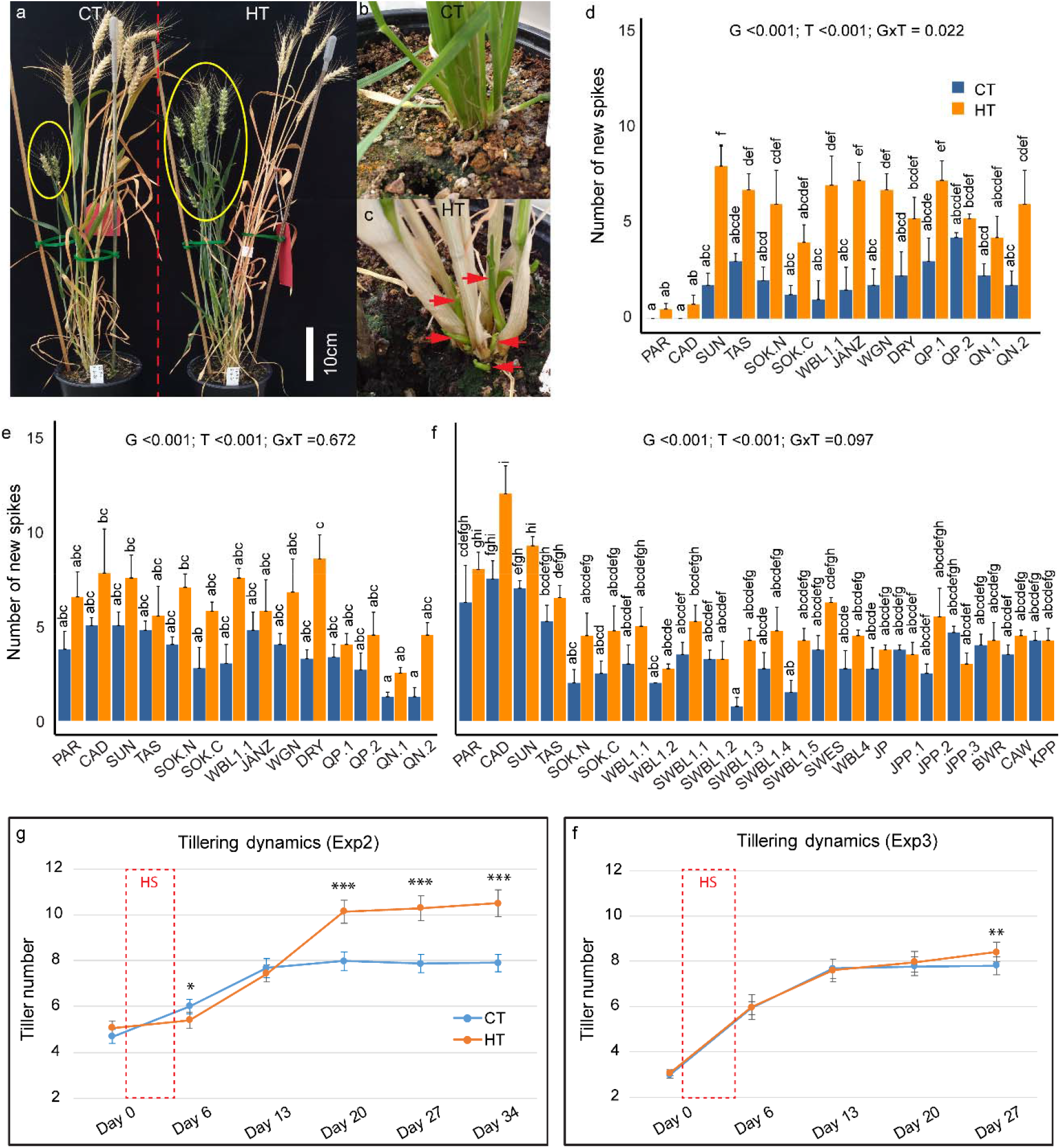
The effects of heat treatments on tillering/spike formation. After heat (HT) treatment, new tillers/spikes (in yellow ellipse) were vigorously stimulated, while control not (CT, a). Tiller outgrowth from CT (b) and HT (c) around one week after stress treatment. A comparison of the number of new spikes between CT and HT in Exp1 (d), Exp2 (e) and Exp3 (f). The effects of genotype/G, treatment/T, interaction/G x T were indicated for each panel. Dynamic change of tillering before (day 0), after (day 6) the 5-day HT treatment (day 1 - 5), and weekly intervals in Exp2 (g) and Exp3 (h). Significance level: P < 0.001 ***; P < 0.01 **.

### Heat stimulated tillering/spike formation and its association with yield

During the ripening stage of Exp1, the senescence status of tillers/spikes was clearly separated into two groups (Fig 2 a) and tillers were therefore distinguished into old (pre-heat) and new (post-heat) spikes for each plant. About one week after heat treatment, new tiller outgrowth was noticed from the bottom of HT stressed plants (Fig 2 b, c). A final count of spikes found significantly more new spikes in HT-treated plants compared to CT plants in Exp1 (p<0.001), Exp2 (p<0.001) and Exp3 (p<0.001) (Fig 2 d, e, f), while the number of old spikes was similar between HT and CT conditions (Fig S5). In addition, there was no significant interaction between treatment and genotype (Fig 2 d, e, f), indicating that all genotypes responded similarly to the HT treatment in terms of new spike formation. The analysis of tillering dynamics in Exp2 and Exp3 showed that onset of new tiller development commenced at two to three weeks after the HT treatment, with a stronger effect observed in Exp2 (Fig 2 g, h). Moreover, the more severe heat stress in Exp2 (37°C/27°C) also caused tiller retardation on day 6, one day after end of the HT treatment (Fig 2 g), but this was not observed under the milder heat stress in Exp3 (34°C/28°C) (Fig 2 h).

As new tillers developed after the HT treatment and extended the days to maturity of the plants, the aboveground biomass per plant (including both old and new tillers) was very similar between HT and CT treatments (Fig 3 a, b, c). Nevertheless, the overall grain yield per plant was significantly reduced after the HT treatment in all the three experiments (p<0.001 for all) (Fig 3 d, e, f). This was primarily due to heat-induced sterility in the old spikes (Fig 3 g, h, i). However, heat-induced formation of new spikes gave rise to similar (Exp2, Fig 3 k) or even significantly higher grain yield from new spikes in Exp1 and Exp3 (Fig 3 j, l). The proportion of yield from new spikes after the HT treatment was therefore significantly higher than under CT conditions (Fig S6). As very limited florets from old spikes set seeds under HT conditions reducing sink size, source supply was more than sufficient and both, width and length of the few developed grains were significantly higher compared to grains from control plants (Table S2). By contrast, the grains from new spikes showed variable responses in terms of width and length (Table S2).

**Fig 3.**
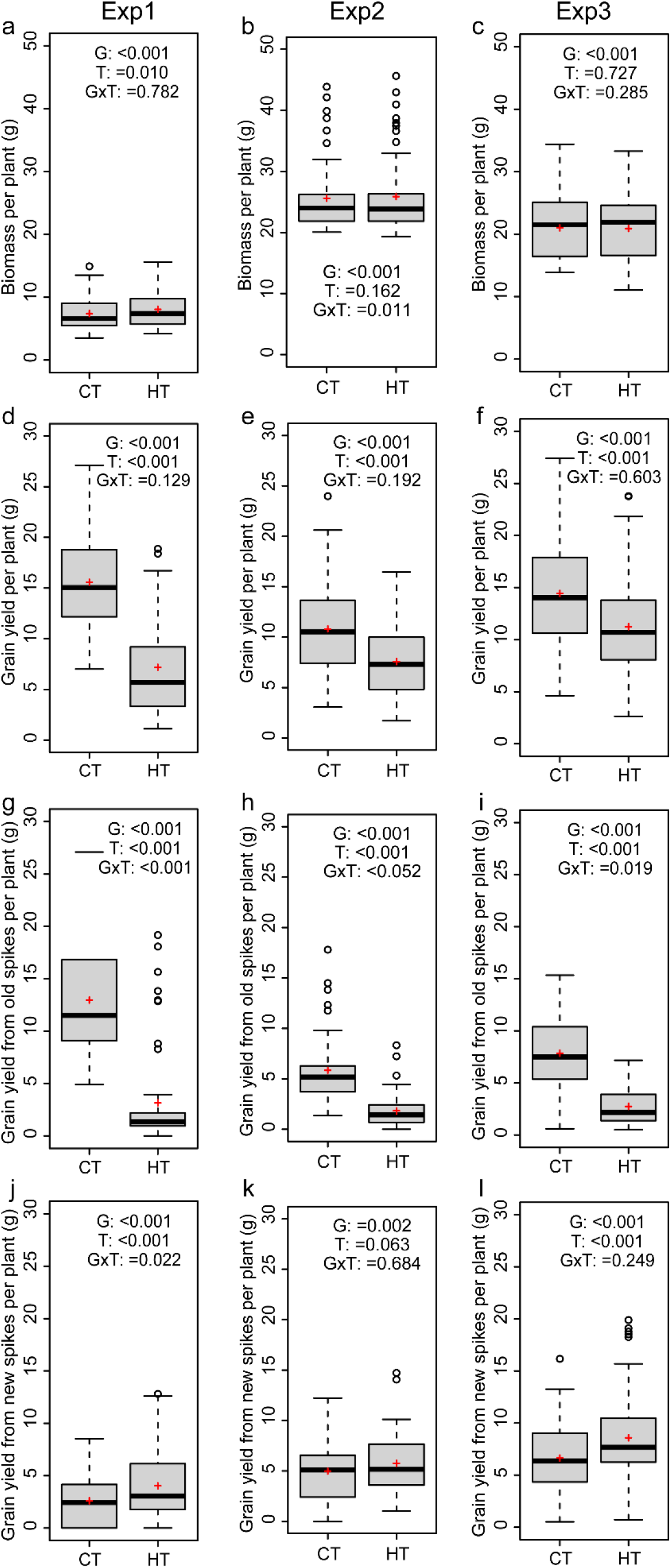
Average values of wheat genotypes grown under high temperature (HT) or control (CT) across the three experiments are shown for biomass (a, b, c), grain yield per plant (d, e, f), grain yield from old spikes (g, h, i), and grain yield from new spikes (j, k, l). The effects of genotype (G), treatment (T), interaction (G x T) are indicated for each panel.

### Heat effects on plant morphology, phenology and chlorophyll dynamics

When wheat plants were exposed to heat stress during the early booting stage, the increase in auricle interval length (AIL) (p < 0.001 for Exp1 and Exp2) and plant height (PH) (p <0.001 Exp2) during the 5-day treatments was significantly reduced by the HT of 37°C/27°C compared to the CT of 21°C/15°C (Fig 4 a, b, d). In contrast, the milder HT of 34°C/28°C in Exp3 only marginally affected AIL (p=0.065) and PH (p=0.279) (Fig 4 c, e). HT treatments also changed plant phenology as indicated by the significantly reduced number of days to heading (DTH) (P <0.001 for Exp2 and Exp3) (Fig 4 f g) and days to maturation (DTM) (p <0.001 for Exp1, Exp2 and Exp3) (Fig 4 h).

**Fig 4.**
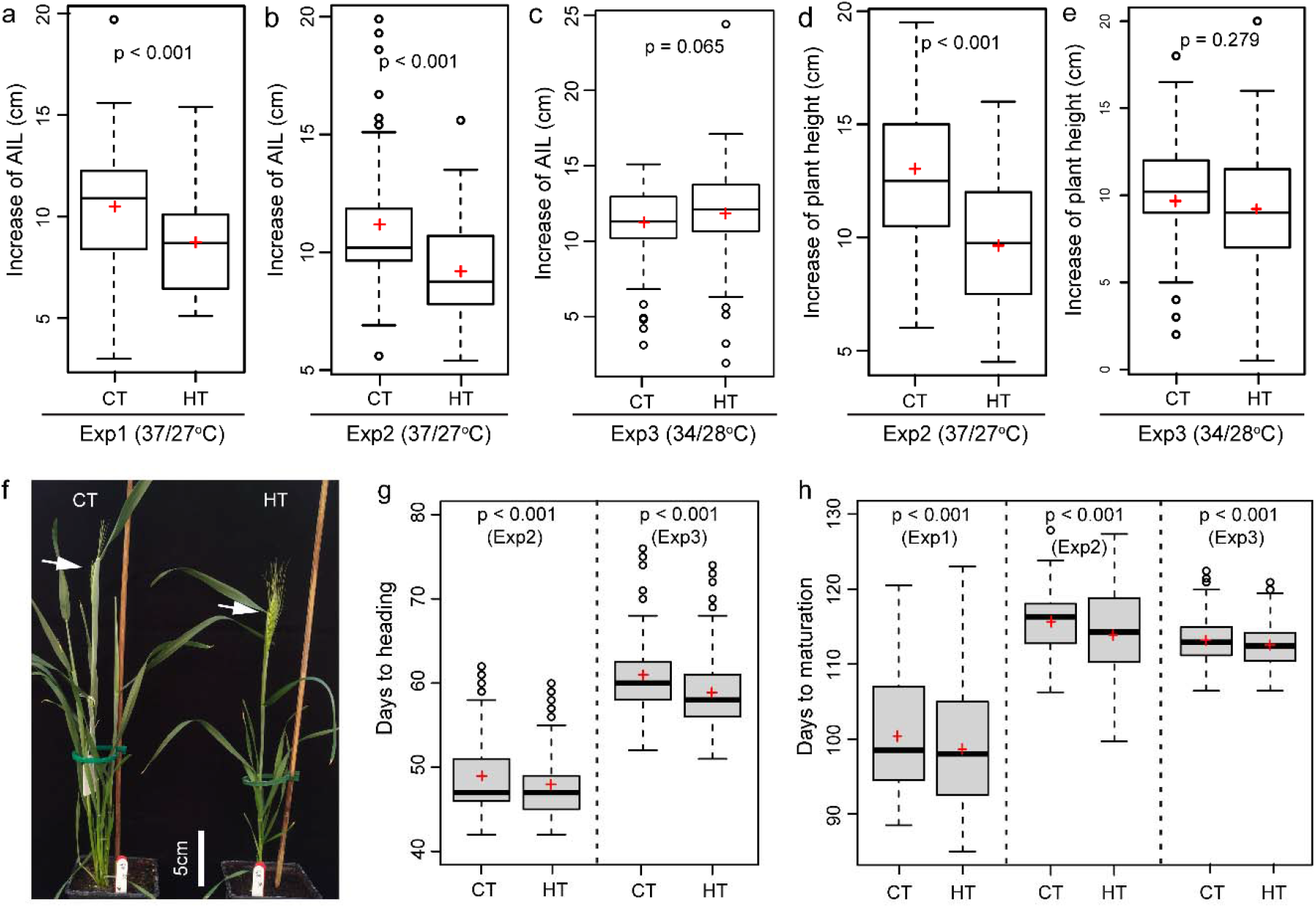
The heat effects on plant morphology and phenology. Comparison of the effect of heat (HT) and control (CT) treatments on the increase of auricle interval length (AIL, a, b, c), the increase in plant height (d, e), days to heading (f, g), and days to maturation (h).

To understand the physiological basis of changes in phenology, dynamic changes in SPAD and Fv/Fm was compared between CT and HT treatments. On day 6 (one day after treatment), in comparison to corresponding CT conditions, SPAD value was significantly reduced by the severe heat (37°C/27°C) in Exp1 (Fig 5 a) and Exp2 (Fig 5 b), but surprisingly increased slightly after the mild heat (34°C/28°C) in Exp3 and maintained a higher maximum SPAD value (Fig 5 e, f). At later stages, however, an accelerated decrease in SPAD was observed under HT conditions in all three experiments, irrespective of heat stress intensity (Fig 5 a, b, e). Based on the time course SPAD measurements, generalized additive models (GAM) were fitted to estimate maximum SPAD (SPADmax), senescence onset (SenOnset) and senescence rate (SenRate) for each combination of genotype and treatment. In Exp1 and Exp3, SenOnset from HT treatment was reproducibly and significantly advanced in comparison with CT conditions, whereas SenRate was similar between treatments (Fig 5 c f). By contrast, Exp2 showed an opposite response with similar SenOnset between treatments, but an increased SenRate under HT (Fig 5 d). This variation between the three experiments may be due to variable intensities of natural sunlight. Even within the same experiment, some genotypes showed earlier SenOnset, while others showed faster SenRate under HT treatment (Fig S7-S9). In addition, the Fv/Fm value at day 6 was always significantly reduced by HT treatment in all three experiments indicating a negative effect of the HT on PSII (Fig S10).

**Fig 5.**
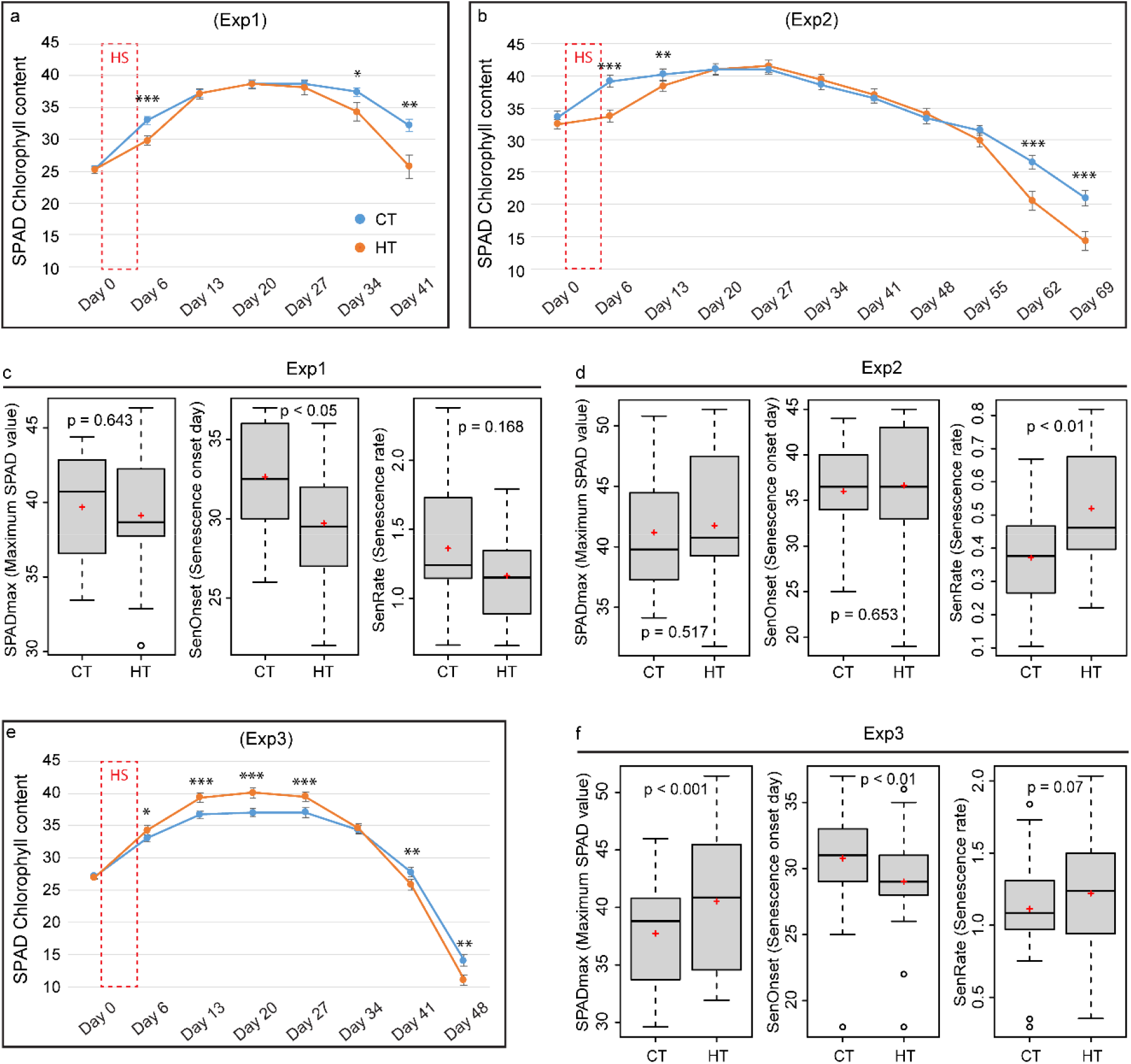
The effects of heat on SPAD (chlorophyll content index) and leaf senescence parameters. Comparison of the effect of heat (HT) and control (CT) treatments on the dynamic change of SPAD measured before (day 0) and after one day after (day 6) a 5-day HT treatment, and in weekly intervals thereafter in Exp1 (a), Exp2 (b) and Exp3 (e). Senescence related parameters, SPADmax (maximum SPAD value), SenOnset (Senescence onset day) and SenRate (Senescence rate), were compared between CT and HT for Exp 1 (c) Exp2 (d) and Exp3 (f). Significance level: P< 0.001 ***; P < 0.01 **; P < 0.05 *.

### Analysis of trait correlations from different experiments and temperature conditions

To understand the relationships among different traits across genotypes, correlations were calculated (Table S3) and visualized as networks (Fig 6). Grain number per spike (GpS) showed different correlations under control and HT conditions; In Exp1 and Exp2, there was no correlation between GpS and any other trait under HT, but under CT, it was positively correlated with spikelet number (SpikeletN) and length (SpikeL) of the tagged spike, as well as with biomass and yield in Exp1. In Exp3, GpS also associated different traits between CT and HT. The importance of induced new tillers and spikes after heat stress was corroborated by the reproducible positive correlations between GY.NT (grain yield of new tillers) and GY (total grain yield per plant), observed in all three experiments (Fig 6, Table S3). This suggests a critical role of new spikes in mitigating heat-induced yield reduction. In addition, the morphological traits, increase in AIL and PH, were generally positively correlated with yield or biomass-related traits, regardless of temperature treatments. Ultimately, SPAD and Fv/Fm were not consistently correlated with other traits from different experiments and treatments. In Exp1, SenOnset showed HT-specific positive correlation with GY.OT (grain yield of old tillers) and SpikeN.NT (spike number of new tillers); SenRate was closely related to yield traits in both Exp2 and Exp3, but not heat-specific; in Exp2, SPADmax was important as it was strongly correlated with several other traits (Fig 6, Table S3).

**Fig 6.**
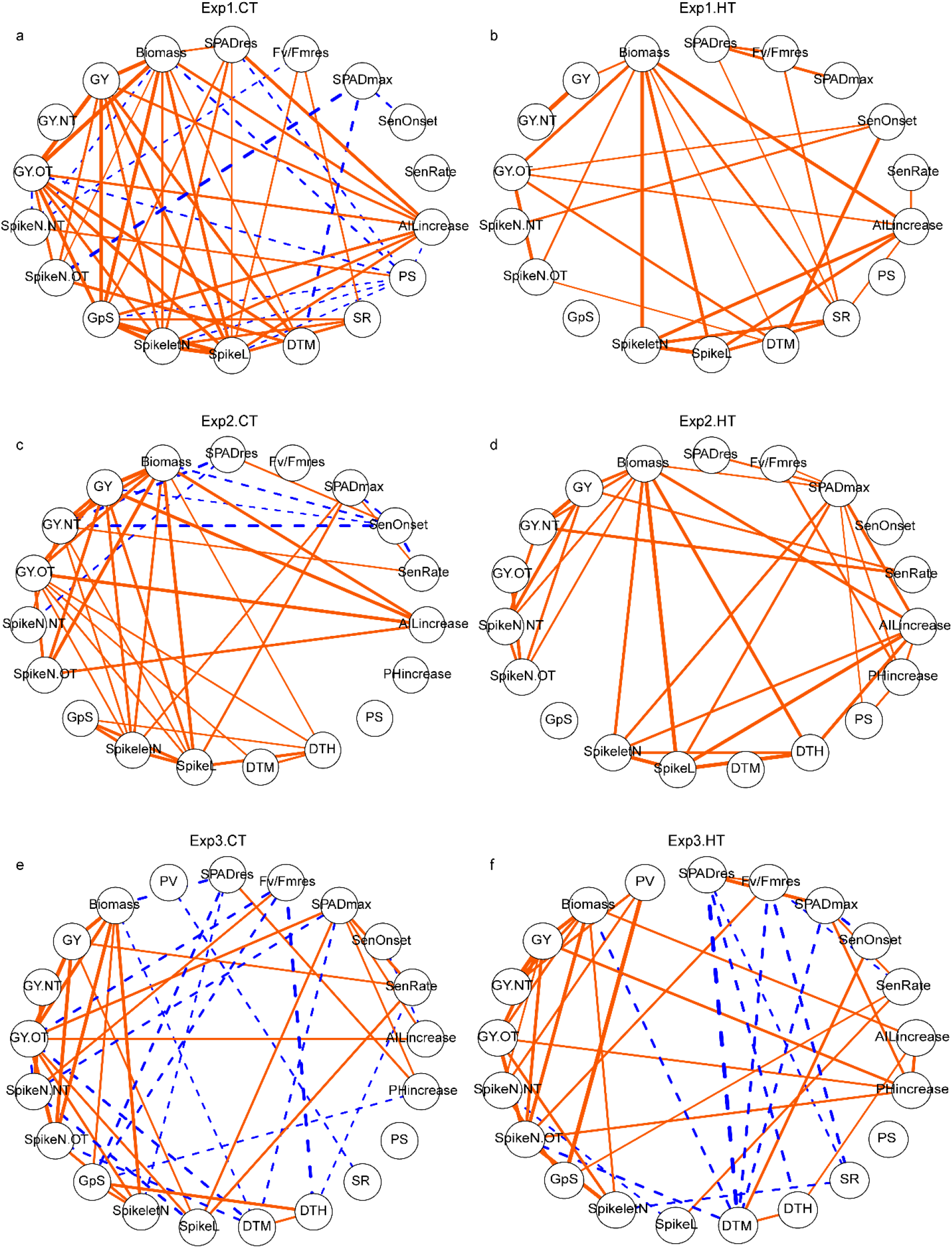
The effects of heat on trait relationships across different experiments. Correlation networks for Exp1 (a, b), Exp2 (c, d) and Exp3 (e, f). Only significant correlations are shown and the edges’ width indicated correlation r value. Orange solid edges represent positive correlations, while blue dashed edges represent for negative correlations. Trait abbreviations: *SPADres*: (SPAD at day 6 – SPAD at day 0)/SPAD at day 0; *Fv/Fmres*: (Fv/Fm at day 6 -Fv/Fm at day 0)/(Fv/Fm at day 0); *SPADmax*: maximum value of SPAD; *SenOnset*: senescence onset time; *SenRate*: senescence rate; *AILincrease*: auricle interval length increase during the 5 day treatment; *PHincrease*: plant height increase during the 5 day treatment; *PS*: observed frequency of paired spikelet from all spikes; *SR*: observed frequency of sham ramification from all spikes; *DTH*: days to heading; *DTM*: days to maturation; *SpikeL*: spike length (tagged primary spike); *SpikeletN*: spikelet number (tagged primary spike); *GpS*: grain number per spike (tagged primary spike); *SpikeN*.*OT*: spike number (old tillers); *SpikeN*.*NT*: spike number (new tillers); *GY*.*OT*: grain yield (old tillers); *GY*.*NT*: grain yield (new tillers); *GY*: grain yield per plant (sum of old and new tillers); *Biomass*: the dry weight of all straw per plant; *PV*: pollen viability from the middle spikelet of tagged spike.

## DISCUSSION

### Importance and limitation of pollen viability as a target trait for wheat heat research

In the present study, anther morphology was gradually affected under two levels of heat stress, 34°C/28°C (day/night) and 37°C/27°C, applied for five days during early booting stage coinciding with pollen development. The more severe heat stress in this study led to a complete loss of pollen viability, while results from a parallel study where the same 37°C/27°C heat treatment lasted for only 3 days (Erena 2018) were similar to the 5-day, milder temperature (34°C/28°C) treatment here, suggesting both stress intensity and duration are critical to screening reproductive heat tolerance. Under the 34°C/28°C condition, pollen viability was considerably variable among genotypes. Two of the lines (SWB1.1 and SWES) with high pollen viability share one common parent, Sokoll, in their pedigree. Sokoll is an advanced wheat line derived from synthetic hexaploid wheat and has shown a yield advantage under terminal heat stress in other reports (Cossani and Reynolds 2015; Thistlethwaite et al. 2020), although it did not show particularly high pollen viability after early booting-stage heat stress in this study. These results suggest stage-specific heat tolerance, therefore, it is necessary to pyramid tolerant traits across different developmental stages. Another parental line included in this study, Weebill1 (WBL1.1 and WBL1.2), has previously been reported to be tolerant to a wide range of variable environmental conditions (Singh et al. 2007). One of the most tolerant genotypes identified in this study was Sunstar, in agreement with data reported by (Erena 2018) who also demonstrated the reproductive heat tolerance of Sunstar. These identified genotypes with heat tolerance during pollen development may be suitable donors for breeding, and warrant further studies to understand the underlying genetic and molecular-physiological mechanisms. The importance of pollen viability is supported by its positive correlation with grain number per spike under heat stress. Interestingly, similar relationships have been reported in other crops (Xu et al. 2017; Shi et al. 2018) and abiotic stresses (Ji et al. 2010), indicating that pollen fertility is a general limiting factor for final grain number under suboptimal growth conditions. Therefore, it should be an important target trait for heat related research and breeding. Nevertheless, the response of pollen viability to heat stress is highly dependent on the developmental stage when stress is applied (Saini and Aspinall 1982; Prasad and Djanaguiraman 2014) and it is thus important to consider genotypic differences and carefully target meiosis to microspore stage when applying heat stress to exclude confounding effects. Currently, the most widely used morphological marker for pollen developmental stage is AIL also known as auricle distance (Ji et al. 2010; Erena 2018; Bokshi et al. 2021). However, AIL corresponding to a specific pollen developmental stage varies among different genotypes (Erena 2018) and must be determined for each genotype, which is laborious. Fortunately, progress has been made by non-destructive X-ray micro computed tomography scanning (Fernández-Gómez et al. 2020) and integrating this with modeling could be a promising way to overcome difficulties with accurate identification of wheat pollen developmental stages.

### Utilizing developmental plasticity to mitigate heat effects on yield

The number of spikes per plant, interacting with spikelet number and floret fertility, determines grain number and thereby final yield. Our data show that a short episode of heat stress during early booting stage induced the development of new tillers and spikes, which is in agreement with other studies (Bányai et al. 2014; Hütsch, Jahn & Schubert 2019; Chavan, Duursma, Tausz & Ghannoum 2019). Though under severe heat stress tillering was initially inhibited, new tillers started emerging at two weeks after recovery, corresponding to about one week after anthesis. This timing suggests that available photosynthates stored in vegetative tissue that cannot be translocated into grain due to spikelet sterility, can be re-allocated into the development of new tillers and spikes. Additional photo-assimilates for new tillers and spikes would be produced during recovery and this is reflected by its positive correlation with delayed onset of senescence (Fig 6 C). The observed formation of new spikes after heat stress compensating for heat-induced biomass and yield losses under controlled environment conditions, now needs to be corroborated under field conditions to ensure that it is a valid target trait for breeding. In addition, a higher frequency of paired spikelets (Boden et al. 2015) and sham ramification (Amagai et al. 2017) was observed in heat treated plants and this may also be related to excessive source supply. Although these traits were not correlated with yield, they could contribute to understand mechanisms underlying such developmental abnormalities.

### Accelerated leaf senescence after brief heat stress during early booting stage

Screening wheat for heat tolerance in the field is generally implemented by late-sowing to impose continuous terminal heat stress during grain filling, often resulting in accelerated leaf senescence (Bergkamp et al. 2018). In the present study, a similar stimulation of flag leaf senescence was observed after the brief heat stress applied during early booting. It is possible that plants are able to measure and memorize phenology or leaf age to program the senescence process (Woo et al. 2019). In our study, model fitting using SPAD time course data proved to be successful in identifying senescence parameters. Both earlier onset and faster senescence rate were identified and were closely related to accelerated leaf senescence, in agreement with similar results reported by (Šebela et al. 2020). Heat-specific positive correlations between senescence onset (SenOnset), new spike formation (SpikeN.NT), and yield from old tillers (GY.OT) in Exp1 supports the important role of late senescence. The observed positive associations between senescence rate (SenRate) and yield traits (grain yield/GY, grain number per spike/GpS, spike length/SpikeL) in Exp2 and Exp3 suggest fast nutrient remobilization in high-yielding lines. Finally, both SPAD and Fv/Fm were reduced by heat immediately after the treatment (day6) in Exp1 and Exp2, but the mild temperature of 34°C/28°C only decreased Fv/Fm, not SPAD. These results indicate Fv/Fm may be more sensitive and therefore a better parameter for heat tolerance evaluation (Cao et al. 2019). Therefore, these senescence-related parameters are useful for crop phenotyping, and integrating modeling with high-throughput imaging measurements will enable large-scale analysis.

## CONCLUSIONS

In this study, a spring wheat panel, including heat tolerant elite varieties and their pre-breeding lines, was dissected for reproductive, developmental, physiological, and yield responses and their inter-relationships after a 5-day heat stress during the early booting stage. In comparison with the control treatment, pollen viability from the tagged primary spike was significantly decreased by heat and subsequently reduced grain number per spike. The heat stress, however, resulted in late tillering after the disruption of sink strength. Consequently, more new spikes were formed contributing to final yield and biomass, though an additional week was needed for the maturation of the late tillers. Flag leaf SPAD (Chlorophyll content index) and Fv/Fm (maximum potential quantum efficiency of Photosystem II) were reduced by heat stress. Model fitting with time course SPAD measurements showed accelerated leaf senescence by either earlier onset or faster senescence rate and these parameters were associated with yield traits. Ongoing genomic and genetic studies will subsequently be used to dissect the mechanism of identified heat-tolerant genotypes (Sunstar, SWBL1.1). Taken together, these reproductive, developmental and physiological traits could be further used as targets for understanding basic mechanisms and breeding heat-tolerant wheat.

## AUTHOR CONTRIBUTIONS

SH conceived and supervised the project, together with EMVS and MJP. JX designed and implemented the experiments and analyzed the data, with support from CL. MPR and SD advised on the selection of genotypes included in this study and provided the seeds. SH, EMVS, SGHL and MJP held regular project planning discussions. JX wrote the manuscript, which was reviewed and edited by SH, CL, MPR, SD, EMVS, SGHL, and MJP.

## ETHICAL STANDARDS

This article does not contain any studies with human participants or animals performed by any of the authors.

## ACKNOWLEDGEMENTS

We would like to thank Tess Rose and Maria Oszvald for their support and suggestions on the manuscript, as well as Matthew Dale for advice on the experimental setup. We would also like to thank Fiona Gilzean, Jill Maple and Jack Turner for taking excellent care of the plants and for their technical support, as well as Chris Hall for helping with the sample processing. Seeds were kindly provided by CIMMYT and Nick Collins, University of Adelaide. This project was funded by the BBSRC UK-Mexico Newton Fund (BB/S012885/1 “Safeguarding Sonora’s Wheat from Climate Change”). SH and MP were also supported by the Designing Future Wheat (DFW) Institute Strategic Programme (BB/P016855/1).

